# RKMR: A Rapid Kernel Machine Regression Framework for Optimal Marker Detection in Spatial Omics Data

**DOI:** 10.64898/2026.07.27.740999

**Authors:** Souvik Seal, Anirban Chakraborty, Chloe Mattila, Mark Rubinstein, Peggi Angel, Debashis Ghosh, Dongjun Chung, Brian Neelon

## Abstract

High-throughput spatial omics technologies enable molecular profiling within intact tissue architecture, yet identifying concise, predictive, and biologically interpretable marker panels for cell types, tissue domains, and disease-associated tissue classes remains challenging. This limitation hinders the development of actionable panels for targeted validation and downstream translation. Existing pipelines rely largely on univariate differential-expression analyses, which ignore joint molecular structure and provide limited predictive insight. Multivariate machine-learning methods, including random forest, XGBoost, elastic net, and specialized single-cell panel-selection approaches, can capture predictive patterns but typically lack explicit spatial modeling and probabilistic feature selection, relying instead on model-specific importance scores or user-specified panel sizes. We develop rapid kernel machine regression (RKMR), a scalable framework for spatial-omics marker discovery that integrates nonlinear kernel modeling, spike-and-slab variable selection, and spatial dependence. RKMR uses automatic relevance determination (ARD) kernels and sparsity-inducing priors to capture nonlinear marker–outcome relationships and implicit feature interactions while producing approximate posterior inclusion probabilities (PIPs) that quantify model-based uncertainty in feature inclusion. To scale inference to large spatial datasets, RKMR combines low-rank kernel approximations with stochastic variational optimization. In simulations, RKMR consistently achieves higher AUPRC than competing methods across a range of molecular-signal and spatial-effect settings. Across spatial transcriptomics and scRNA-seq datasets, RKMR identifies parsimonious marker sets that recover reported cell-type signatures and reproducible tissue-layer markers. These results establish RKMR as a scalable and uncertainty-aware framework for translating high-dimensional spatial omics data into robust, experimentally actionable marker panels.

## 1 Introduction

The FDA–NIH *Biomarkers, EndpointS, and other Tools (BEST)* resource defines a biomarker as a measurable characteristic that reflects a biological process, disease state, or response to an intervention^1^. In bulk and single-cell omics, biomarkers include DNA mutations, transcriptional signatures, and protein abundance patterns used for disease classification, tissue and cell-state annotation, and therapeutic decision-making ^2–5^. Spatial transcriptomics (ST) and spatial proteomics (SP) expand this concept by enabling discovery of disease-relevant molecular signals that vary across spatially localized cell populations, tissue domains, and disease states ^6–9^. For clinical translation, however, markers must be not only discriminative but also compact and maximally informative. Established assays such as Oncotype DX (21-gene recurrence score) and EndoPredict (12-gene molecular score) demonstrate how compact molecular panels can support treatment decision-making in early-stage breast cancer^10–13^. As ST and SP technologies become increasingly scalable, they offer opportunities to translate tissue-resolved biology into targeted molecular panels for precision medicine ^14–17^. Despite this promise, many spatial discovery workflows in ST remain centered on spatially variable gene (SVG) or cell-type-specific SVG detection ^18–23^. These approaches are useful for exploratory identification of genes whose expression varies across tissue space or within spatially localized cell populations, but spatial variation alone does not ensure predictive discrimination, parsimony, or quantified evidence for inclusion in a targeted marker panel. The present work addresses this gap by focusing on predictive marker discovery while accounting for redundancy, uncertainty, and spatial organization.

In the supervised marker-discovery setting, outcome annotations, such as reference-derived cell types or expert-annotated tissue layers, are assumed to be known. We focus on binary *one-versus-rest* comparisons, where the goal is to discriminate a target class from all remaining classes. Most marker-detection workflows for bulk and single-cell RNA sequencing (scRNA-seq) data still rely on marginal differential expression (DE) tests, in which features are evaluated individually and ranked by effect size or statistical significance ^24–32^. Compact panels are then derived through ad hoc filtering of top-ranked candidates rather than by explicitly modeling joint predictive value, redundancy, and selection uncertainty^33^. Growing evidence suggests that such approaches can suffer from limited reproducibility across datasets and analytical settings ^34^. More recent work has shifted toward compact marker-panel selection using supervised, multivariate, combinatorial, or constrained optimization frameworks, including COMET ^35^, RANKCORR^36^, scGeneFit ^37^, NS-Forest ^38^, COSG^39^, SMaSH ^40^, MAGNETO^41^, CellBRF ^42^, MarkerMap ^43^, Spapros ^44^, scGIST ^45^, FSCME^46^, and gps-FISH^47^. These methods represent important progress, but they differ substantially in their objectives, as summarized in Table 1, and were developed primarily for scRNA-seq datasets, which may limit their performance and interpretability in spatial omics applications. Moreover, most lack formal uncertainty estimates for marker inclusion, depend on user-specified panel sizes rather than adaptive selection criteria, and scale poorly with increasing sample size. Together, these limitations motivate automated, scalable, and spatially informed methods for marker discovery.

**Table 1:** Summary of specialized marker-selection methods categorized into two broad classes: *one-vs-rest* and *multiclass*. Methods shown in italics were included in our benchmark because their software packages are readily usable and compatible with Apple M-series Macs and recent versions of `Python`.

| Selection scope | Method | Test type | Basic algorithm | Selection output |
| --- | --- | --- | --- | --- |
| One-versus-rest | COMET<sup>35</sup> | Nonparametric testing | Applies expression thresholding and hypergeometric testing of small gene combinations. | Deterministic significance-based ranking; final panel is user-selected. |
|  | RANKCORR<sup>36</sup> | Rank-based optimization | Rank-transforms expression and performs sparse optimization. | Deterministic and non-probabilistic; panel size is controlled by a user-specified sparsity parameter. |
|  | NS-Forest<sup>38</sup> | Repurposed random forest (RF) | Combines RF importance, expression thresholds, and combinatorial scoring. | Non-probabilistic but stochastic; panel size is selected algorithmically. |
|  | *COSG*<sup>39</sup> | Cosine-similarity scoring | Ranks genes by cosine similarity with the target cell-type expression profile. | Deterministic and non-probabilistic; the number of retained genes is user-defined. |
|  | SMaSH<sup>40</sup> | Nonlinear machine learning | Fits ensemble or neural-network classifiers and ranks genes using feature-importance scores. | Non-probabilistic but stochastic; the number of retained markers is user-defined. |
|  | *MAGNETO*<sup>41</sup> | Evolutionary optimization | Optimizes classification performance and panel size using a bi-objective evolutionary algorithm. | Non-probabilistic but stochastic; the final Pareto-optimal panel is user-selected. |
| Multiclass | *scGeneFit*<sup>37</sup> | Sparse multiclass optimization | Jointly selects genes that preserve separation among multiple cell types. | Deterministic and non-probabilistic; target panel size is user-defined. |
|  | CellBRF<sup>42</sup> | Repurposed RF | Generates cluster-based pseudo-labels and selects genes using RF importance. | Non-probabilistic but stochastic; panel size is determined by an importance threshold. |
|  | MarkerMap<sup>43</sup> | Neural-network feature selection | Learns a fixed-size gene subset using neural-network-based selection. | Stochastic differentiable feature selection using Gumbel-Softmax; panel size is user-defined. |
|  | Spapros<sup>44</sup> | Sequential multi-criterion selection | Sequentially combines variation, discrimination, differential expression, and probe-design criteria. | Non-probabilistic sequential selection; panel size and prior-gene constraints are user-defined. |
|  | scGIST<sup>45</sup> | Constrained neural-network selection | Uses a sparse feature-selection layer with penalties for panel size and prioritized genes. | Non-probabilistic but stochastic; panel size and gene priorities are user-defined. |
|  | FSCME<sup>46</sup> | Entropy-weighted correlation filtering | Combines copula correlation and maximal information coefficient using entropy-derived weights. | Deterministic and non-probabilistic; subset size or selection cutoff is user-defined. |
|  | *gpsFISH*<sup>47</sup> | Genetic algorithm | Evolves candidate panels using cross-validated classification performance as fitness. | Non-probabilistic but stochastic; panel size is user-defined. |

Kernel machine regression (KMR) became prominent in genome-wide association studies (GWAS) for detecting nonlinear associations between genetic variants and complex phenotypes ^48–54^. KMR models feature effects through a semiparametric function, often represented as a Gaussian process (GP), whose kernel covariance encodes sample similarity based on observed features ^55^. Recent kernel-based approaches have extended differential expression analysis in single-cell data beyond mean shifts to nonlinear distributional differences ^30,31^. However, these methods remain focused largely on individual gene-level testing rather than predictive marker-panel selection, typically constructing kernels from one gene at a time. Moving from testing to marker selection requires kernels that encode feature-specific relevance across many candidate markers. Automatic relevance determination (ARD) kernels provide this mechanism by allowing features to contribute unequally within the kernel framework ^56–60^. When combined with variable-selection priors, such as spike-and-slab priors ^61^, ARD can shrink uninformative features toward zero and support adaptive marker selection. This motivates a Bayesian logistic KMR (BLKMR) model in which known cell or tissue annotations serve as the outcome, molecular features are embedded within an ARD kernel, and marker evidence is summarized by posterior inclusion probabilities (PIPs) ^62,63^. In spatial omics, however, these annotations are observed within intact tissue architecture, where nearby spots or cells often exhibit spatial autocorrelation—that is, similar cell types or tissue states tend to cluster in physical space because of shared microenvironmental context^64–66^. A spatially augmented BLKMR framework can therefore jointly model marker relevance and spatial dependence.

A Bayesian Markov chain Monte Carlo (MCMC) implementation of such a spatial BLKMR model is computationally prohibitive, as dense matrix operations require *O*(*N* ^3^ + *Np*^2^) floating-point operations (FLOPs) per update, where *N* denotes the number of spots or cells and *p* denotes the number of molecular features ^67^. This burden is especially limiting for modern spatial omics datasets, where *N* can approach or exceed 10^5^ and *p* can exceed 10^4^. Recent work by Dance and Paige (2022) ^68^ developed a highly scalable algorithm for ARD-based KMR with continuous outcomes, using approximate coordinate-ascent variational inference and minibatching ^69,70^. Building on this strategy, we develop an approximate spatial BLKMR algorithm based on a penalized quasi-likelihood (PQL) approach, accelerating key intermediate updates through stochastic variational inference and Nyström kernel approximation ^71–73^. These approximations drastically reduce the dominant computational burden, with empirical timing studies showing near-linear scaling in both *N* and *p*. We refer to the resulting framework as rapid kernel machine regression (RKMR). In complex simulation scenarios, RKMR achieves superior marker-selection accuracy and specificity compared with specialized marker-selection methods and general-purpose algorithms, including random forest, XGBoost, and elastic net, with area under the precision-recall curve (AUPRC) improvements of up to 50% in several settings. Across multiple scRNA-seq and ST datasets on diverse cancers and anatomical structures, RKMR reliably recovers expert-annotated markers of distinct cell types and tissue layers, while alternative methods exhibit less stable recovery. Together, these results position RKMR as a practical and scalable probabilistic framework for targeted marker-panel discovery, supported by an efficient Python implementation for broad use in single-cell and spatial omics studies.

## 2 Methods

KMR offers a flexible semiparametric framework for capturing nonlinear feature effects through kernel covariance functions and is closely connected to GP regression ^74^. Building on its long-standing development in statistics and machine learning, including Bayesian extensions for regression, classification, and variable selection ^75–78^, we develop a spatially augmented, ARD-based BLKMR model for marker discovery. Pairing it with scalable approximations, we term the framework rapid KMR (RKMR). We first describe the model formulation, then introduce sparsity-inducing priors for feature selection, and finally present the scalable inference algorithm. We also briefly review existing specialized marker-selection methods (Table 1).

### 2.1 RKMR: Rapid kernel machine regression

#### 2.1.1 Model formulation

Consider a *one-vs-rest* classification framework for a given cell type or tissue annotation, where the observational unit may be either a single cell or a spatial spot/location, depending on the platform. For *i* = 1, … , *N* , let *y*_*i*_ denote the binary (0*/*1) label indicating whether the *i*^th^ unit belongs to the target class. For each unit, we consider two sets of covariates: **x**_*i*_ ∈ ℝ^*b*^, which are assumed to have a linear effect on *y*_*i*_ (e.g., batch indicators or other adjustment covariates), and **z**_*i*_ ∈ ℝ^*p*^, which are allowed to have a potentially nonlinear effect on *y*_*i*_ (e.g., gene, protein, or peptide abundance profiles). **s**_*i*_ ∈ ℝ^2^ denotes the location coordinate of unit *i*. We assume *y*_*i*_ *∼* Bernoulli(*µ*_*i*_), where *µ*_*i*_ = Pr(*y*_*i*_ = 1 | **x**_*i*_, **z**_*i*_, **s**_*i*_) denotes the probability that the *i*^th^ unit belongs to the target class. We model *µ*_*i*_ using a canonical logit link:

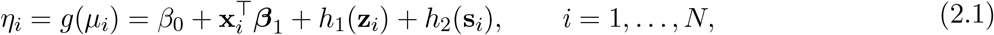

where *g*(·) = logit(·), *β*_0_ is the intercept, and ***β***_1_ is the coefficient vector for the linear covariates **x**_*i*_. The function *h*_1_ : ℝ^*p*^ → ℝ captures the nonlinear effects of **z**_*i*_ and is assumed to lie in a reproducing kernel Hilbert space (RKHS) *ℋ*_*K*_ induced by a positive semidefinite kernel *K*_**r**_(·, ·). Equivalently, we place a mean-zero GP prior on *h*_1_: *h*_1_(·) *∼ GP*(0, *τ*_1_ *K*_**r**_(·, ·)) , where *τ*_1_ *>* 0 is a marginal variance or kernel amplitude parameter. Although *K*_**r**_ can take several forms, we use an ARD radial basis function (RBF) kernel ^57^, where **r** = (*r*_1_, … , *r*_*p*_)^*⊤*^ contains positive feature-specific relevance parameters. Under this prior, the latent vector **h**_1_ = (*h*_1_(**z**_1_), … , *h*_1_(**z**_*N*_))^*⊤*^ satisfies

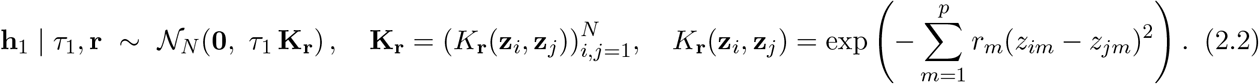

The feature-specific relevance parameter *r*_*m*_ controls how strongly feature *m* contributes to pairwise kernel similarity. Values of *r*_*m*_ close to zero effectively downweight feature *m*, whereas larger values allow differences in feature *m* to more strongly influence the nonlinear component *h*_1_(·) and, consequently, the outcome probability. A simpler alternative is a linear ARD kernel of the form 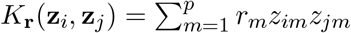, where *r*_*m*_ *≥* 0 controls the contribution of feature *m* to the weighted inner product. Unlike the nonlinear ARD kernel, however, this formulation captures only linear associations between the molecular features and the outcome on the scale of the linear predictor.

Similarly, *h*_2_(·) captures residual spatial autocorrelation in the outcome and is modeled as a latent spatial GP over locations ^64^. Specifically, we assume *h*_2_(·) *∼ GP*(0, *τ*_2_ *C*_*ρ*_(·, ·)) , where *τ*_2_ *>* 0 is spatial variance and *C*_*ρ*_(·, ·) is a valid spatial correlation function indexed by the hyperparameter *ρ*. Consequently, **h**_2_ = (*h*_2_(**s**_1_), … , *h*_2_(**s**_*N*_))^*⊤*^ satisfies

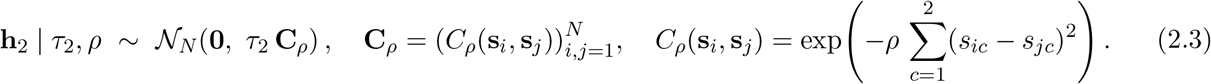

Here, *ρ ≥* 0 is an inverse length-scale parameter; a larger *ρ* implies faster decay of spatial correlation with distance (more local dependence), whereas a smaller *ρ* implies slower decay (longer-range dependence).

#### 2.1.2 Sparsity prior on the relevance parameters

In Bayesian linear models, sparsity-inducing priors, such as the discrete *spike-and-slab* prior or the continuous *horseshoe* prior, shrink negligible effect-size estimates toward zero, thereby enabling selection of the most important features ^79–81^. Because the feature-specific relevance parameter *r*_*m*_ plays an analogous role in the ARD kernel, sparsity-inducing priors can similarly be used to identify relevant molecular features. For computational tractability, we use a continuous approximation to the discrete spike-and-slab prior, replacing the Dirac spike at zero with a Gaussian spike having small variance. Similar continuous spike-and-slab approximations have been used previously in linear regression and GP modeling^82,83,68^. The prior has the following form

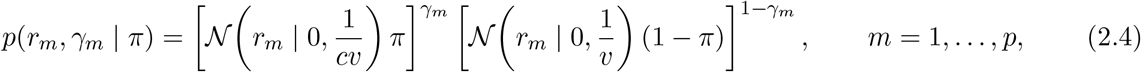

where ***γ*** = (*γ*_1_, … , *γ*_*p*_)^*⊤*^ *∈ {*0, 1*}*^*p*^ are binary inclusion indicators. The constant *v ≫* 1 defines a *spike* distribution with variance 1*/v* concentrated near zero, whereas *cv ≪* 1 defines a more diffuse *slab* distribution with variance 1*/*(*cv*). We further place a Beta prior on the overall inclusion probability *π, p*(*π*) = Beta(*π* | *a, b*). The posterior inclusion probability (PIP) for feature *m* is defined as Pr(*γ*_*m*_ = 1 | data) and is approximated by the variational inclusion probability *λ*_*m*_ as discussed later.

### 2.1.3 Estimation details

The RKMR inference algorithm embeds variational inference (VI) within an outer PQL and iteratively reweighted least squares (IRLS) procedure for the BLKMR model^73,69,70^. At each outer iteration, the Bernoulli likelihood is locally approximated by a weighted Gaussian working model. Suppressing the outer-iteration index for notational simplicity, let *η*_*i*_ denote the current linear predictor from Eq. 2.1, and define *µ*_*i*_ = *σ*(*η*_*i*_), *w*_*i*_ = *µ*_*i*_(1 *− µ*_*i*_), where *σ*(·) denotes the expit, or inverse-logit, function. A first-order Taylor expansion of the inverse-link mean function around the current linear predictor yields the IRLS pseudo-response

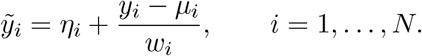

Following Ge et al. ^84^, this produces the local Gaussian working model

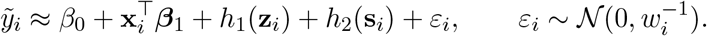

Let **W** = diag(*w*_1_, … , *w*_*N*_). After integrating out the Gaussian random effects *h*_1_(·) and *h*_2_(·), the pseudoresponse has the approximate marginal distribution

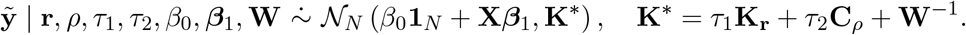

Thus, at each outer PQL/IRLS iteration, the original Bernoulli KMR model is replaced by a local Gaussian surrogate whose pseudo-response and covariance depend on the current fitted probabilities. Rather than applying VI directly to the original non-Gaussian (Bernoulli) mixed model, as in several variational GLMM approaches ^85,86^, RKMR performs VI conditional on the current Gaussian working response 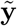 and weight matrix **W**. This formulation reduces inference within each outer iteration to a weighted Gaussian problem, thereby facilitating the use of low-rank Nyström approximations, Woodbury identity-based linear solves, and stochastic-gradient optimization. Under the continuous spike-and-slab prior specified in Eq. 2.4, we approximate the conditional posterior distribution of the ARD relevance parameters **r**, binary inclusion indicators ***γ***, and overall inclusion probability *π* using the mean-field factorization^70^

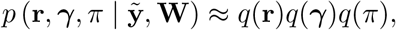

where *q*(·) denotes the corresponding variational posterior distributions. We parameterize the discrete variational factor as 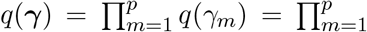 Bernoulli(*γ*_*m*_; *λ*_*m*_),, where *λ*_*m*_ = *q*(*γ*_*m*_ = 1) denotes the variational inclusion probability for feature *m*. We use a Beta variational family for the baseline inclusion probability, *q*(*π*) = Beta(*π*; *ξ*_*a*_, *ξ*_*b*_). Conditional on the current Gaussian surrogate, the variational distributions are estimated by maximizing the evidence lower bound (ELBO)

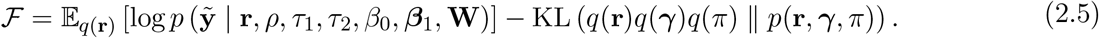

Using the hierarchical factorization *p*(**r**, *γ, π*) = *p*(**r** | *γ*)*p*(*γ* | *π*)*p*(*π*), the Kullback-Leibler (KL) contribution to the ELBO can be decomposed into the expected log-prior and variational entropy terms,

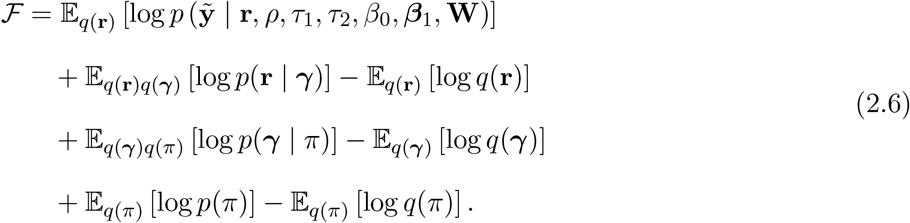

The variational optimization is therefore embedded within the outer PQL/IRLS loop: the ELBO is optimized conditional on the current 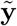 and **W**, after which the fitted linear predictor, working response, and weights are updated. Following Dance and Paige (2022) ^68^, optimization of the surrogate ELBO combines coordinate-ascent and stochastic-gradient variational updates. To improve scalability, the stochastic gradients for the ARD relevance parameters, spatial range parameter, and kernel amplitudes are estimated using randomly sampled minibatches of observations. In contrast, the full-data PQL/IRLS update uses a Nyström approximation to the combined molecular and spatial kernel matrix.

When the second moments 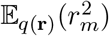 are available in closed form, the conditional conjugacy of the spike-and-slab hierarchy yields closed-form coordinate-ascent updates for *q*(*γ*) and *q*(*π*), conditional on the current *q*(**r**). For each *m*,

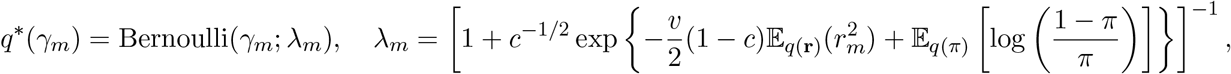

where *c* and *v* are the fixed spike-and-slab hyperparameters defined in Eq. 2.4. The variational distribution of the inclusion probability has the conjugate update

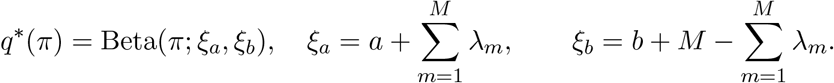

These updates constitute the approximate coordinate-ascent VI (a-CAVI) component of RKMR. The resulting *λ*_*m*_ values quantify the variational posterior probabilities that the corresponding ARD relevance parameters arise from the slab component and can therefore be interpreted as approximate PIPs. In contrast, closed-form coordinate-ascent updates are unavailable for *q*(**r**) because the kernel covariance **K**_**r**_ depends nonlinearly on the ARD relevance parameters. Following Dance and Paige (2022), we impose the zero-temperature restriction 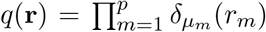 where 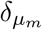 denotes a Dirac point mass concentrated at *µ*_*m*_. Consequently, 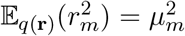 . The mean parameters ***µ*** = (*µ*_1_, … , *µ*_*p*_)^*⊤*^ are then optimized using stochastic gradients of the variational objective (see the Supplementary Material). The spatial correlation range parameter *ρ* and kernel amplitudes (*τ*_1_, *τ*_2_) are similarly updated using stochastic gradients.

Overall, the RKMR algorithm can be summarized as follows. Conditional on the current Gaussian surrogate, stochastic-gradient updates are used for the kernel parameters, while coordinate-ascent updates are used for the inclusion indicators and mixing probability. The linear predictor is then recalculated, yielding updated fitted probabilities, IRLS weights, and pseudo-responses for the next iteration. This process is repeated until convergence, after which the final inclusion probabilities *λ*_*m*_ are used for marker selection. In the Supplementary Material, the full procedure is summarized in Algorithm 1; the Nyströmaccelerated IRLS update is described in Algorithm 2, and the ARD-gradient and inclusion-probability updates are provided in Algorithm 3.

#### 2.1.4 Hyperparameter tuning and PIP thresholding

RKMR includes two hyperparameters, *c* and *v*, that control the relative variances of the Gaussian spike-and-slab prior. Specifically, *v* is the spike precision and *cv* is the slab precision, corresponding to variances 1*/v* and 1*/*(*cv*). As discussed by Dance and Paige (2022), directly optimizing *v* or introducing a variational distribution *q*(*v*) can yield results that are sensitive to initialization. We therefore fixed *c* = 10^*−*8^, ensuring that the spike remained tightly concentrated near zero relative to the slab, and fitted RKMR using 6 fixed candidate values of *v* between 100 and 10^6^. The resulting posterior quantities were combined through model averaging across the candidate values ^87^. Because the models for different values of *v* can be fitted independently, these computations can be parallelized, reducing runtime. The number of Nyström landmarks was fixed at 500. Stochastic-gradient updates were computed using random minibatches of 256 observations. In all real-data analyses, genes with PIP *>* 0.5 were selected as markers. In most cases, the highest-ranked genes had PIPs close to 1, indicating strong posterior support for their inclusion.

### 2.2 A brief review of marker selection methods

Substantial methodological advances have extended marker selection beyond standard univariate differential-expression tests in single-cell data. Table 1 summarizes representative approaches in two broad classes: *one-versus-rest* methods, which identify markers that distinguish a target cell type from all remaining cell types, and *multiclass* methods, which seek marker panels that jointly discriminate among all cell types. Most of these methods are developed primarily for scRNA-seq data, whereas dedicated marker-selection approaches for ST and SP data remain limited. These methods should be distinguished from spatially variable gene-detection approaches, which identify spatial expression patterns but do not directly yield actionable discriminative marker panels. We refer readers to recent developments in spatial marker selection ^88–90^.

## 3 Simulation Study

We consider two simulation scenarios: (1) linear effects of the features on the outcome, generated using a linear ARD kernel, and (2) nonlinear effects, generated using an ARD RBF kernel. In both scenarios, only three features are truly associated with the outcome. Each active feature belongs to one of three correlated blocks of size five, with a common pairwise correlation of 0.5 within each block, reflecting the correlated expression patterns commonly observed among genes. Specifically, we generate the feature matrix **Z** = (**z**_1_, … , **z**_*N*_)^*⊤*^ from a multivariate normal distribution with mean zero and a block-diagonal correlation matrix. The first three blocks follow compound-symmetric correlation structures, with all within-block off-diagonal correlations equal to 0.5, whereas the remaining (*p −* 15) features are mutually independent. This design produces a realistic feature-correlation structure while ensuring that the true signal is confined to a small subset of correlated features. For simplicity, we do not include any additional covariates **x**_*i*_. For each observation, the two-dimensional spatial coordinate **s**_*i*_ is generated by independently sampling each coordinate from a Unif(0, 1) distribution. Finally, the binary outcome *y*_*i*_ is generated according to the KMR model defined in Equations 2.1, 2.2, and 2.3, under varying values of *τ*_1_, *τ*_2_, and *r*_*m*_. Here, *τ*_1_ controls the overall magnitude of the molecular kernel contribution and can therefore be interpreted as the signal strength, whereas *τ*_2_ controls the magnitude of the spatial kernel contribution. For the active features, the relevance parameter *r*_*m*_ is varied over 0.1, 0.5, 1, 2, with larger values corresponding to a stronger influence on the molecular kernel. The spatial range parameter *ρ* is fixed at 0.5 throughout the simulations. The outcome generation mechanisms for the two simulation scenarios are summarized below.

**Scenario 1**. We simulate ***η*** = (*η*_1_, … , *η*_*N*_) as

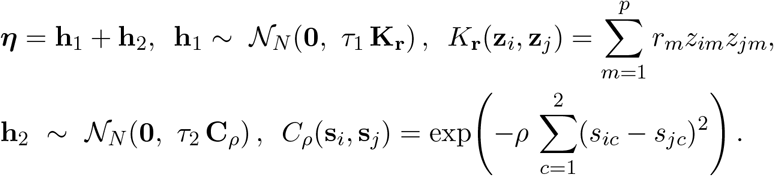

Conditional on *η*_*i*_, the binary outcome is generated as *y*_*i*_ *∼* Bernoulli *{*expit(*η*_*i*_)*}* . In this scenario, the molecular effect is generated using a linear ARD kernel. We vary *τ*_1_ over 5, 10 and *τ*_2_ over 100, 400, 900. Thus, larger values of *τ*_1_ correspond to a stronger overall molecular signal, larger values of *τ*_2_ correspond to a greater contribution of the spatial random effect to the linear predictor, and larger values of *r*_*m*_ correspond to a stronger contribution of feature *m* to the molecular kernel.

**Scenario 2**. We simulate ***η*** as

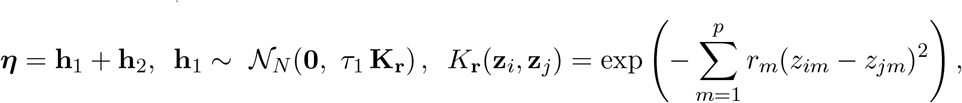

The spatial effect **h**_2_ and binary outcome *y*_*i*_ are generated as in Scenario 1, whereas the molecular effect is nonlinear. We vary *τ*_1_ over 10,50 and *τ*_2_ over 5,10. In comparison to the earlier scenario, we consider larger values of *τ*_1_ and smaller values of *τ*_2_ because identifying the active molecular features is substantially more difficult under the nonlinear ARD RBF kernel, particularly when the spatial effect is strong relative to the molecular signal.

We compare RKMR with general-purpose methods—random forest (RF), XGBoost (XGB), and elastic net (Enet)—and specialized marker-selection methods, including scGeneFit and gpsFISH with panel sizes of 3 and 10, COSG, and MAGNETO. Some of these methods were developed with broader analytical objectives or downstream applications beyond marker recovery; accordingly, our benchmark evaluates only their feature-selection performance in the present setting. Because methods such as SMaSH ^40^ and CellBRF ^42^ build on general-purpose algorithms such as RF and XGB, our comparisons also indirectly assess their underlying learning algorithms. Performance is evaluated using the AUPRC, averaged over 50 replications. For RKMR, precision–recall curves are obtained by varying the PIP threshold over [0, 1]; for competing methods, we use feature-importance scores when available and binary inclusion indicators otherwise. AUPRC is computed using `scikit-learn`.

### 3.1 Simulation Scenario 1

Figure 1 shows that RKMR achieved the highest AUPRC across all parameter settings, indicating consistently strong feature-recovery performance. Within each subfigure, AUPRC increased with *r*_*m*_, as larger values correspond to stronger contributions from the active molecular features and therefore facilitate their separation from inactive features. In contrast, increasing *τ*_2_ from the leftmost to the rightmost subfigure strengthened the spatial effect relative to the molecular signal, making active-feature recovery more difficult and reducing AUPRC across methods. Nevertheless, RKMR maintained its performance advantage throughout. Because this scenario uses a linear ARD kernel and assumes linear gene–outcome associations, Enet also performed competitively and approached RKMR in several settings. Among the specialized methods, scGeneFit and COSG achieved moderate performance but were often outperformed by standard RF and Enet, suggesting limited benefit from their specialized marker-selection strategies in this setting. We provide more results in the Supplementary Material with varying values of *N* and *p*.

**Figure 1:**
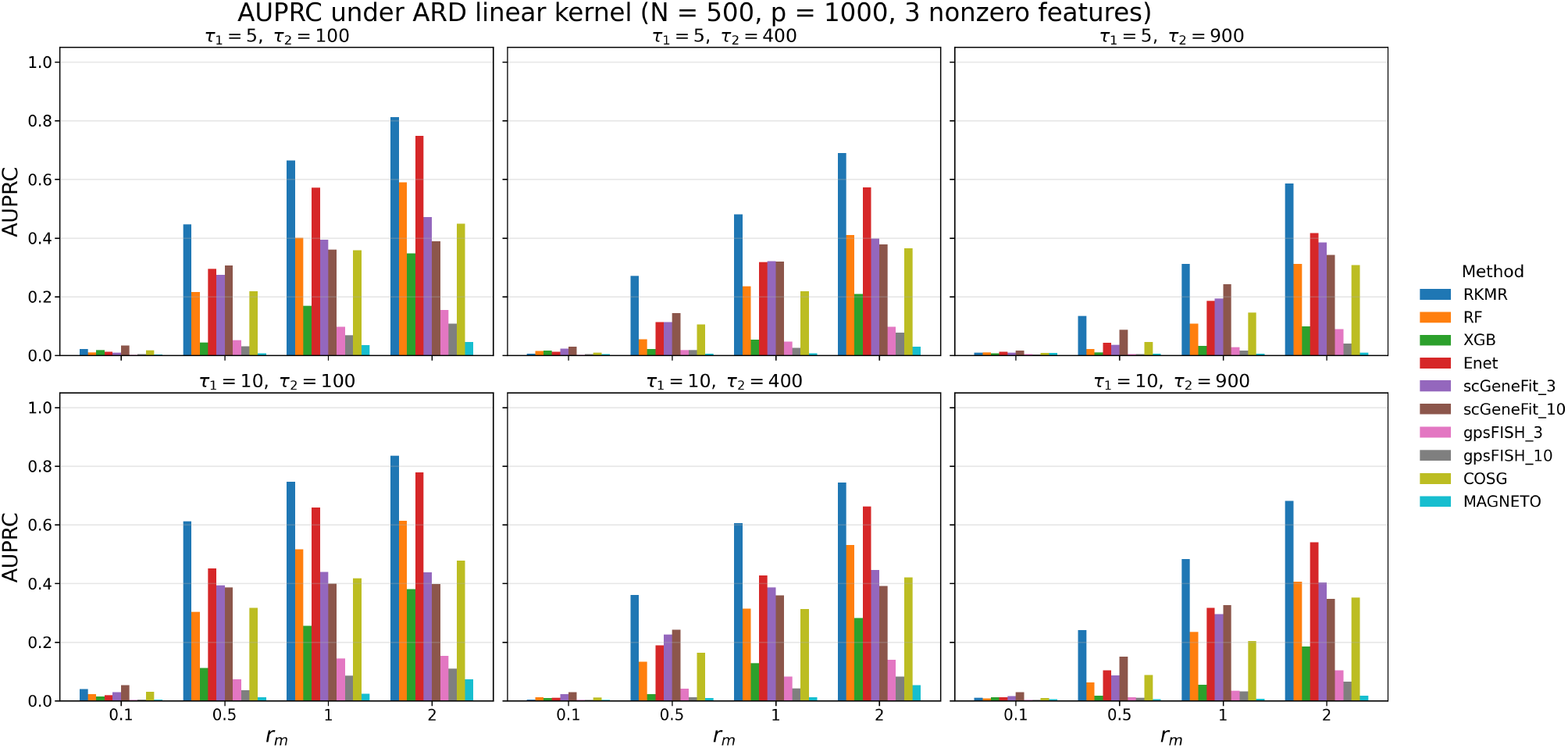
AUPRC comparisons for Simulation Scenario 1 across parameter settings. Within each subfigure, increasing *r*_*m*_ corresponds to greater relevance of the active features. From left to right, the spatial effect increases relative to the molecular effect, making active-feature recovery more challenging.

### 3.2 Simulation Scenario 2

Feature relevance under an ARD RBF kernel requires careful interpretation. A larger *r*_*m*_ increases the contribution of feature *m* to the kernel distance. However, when *r*_*m*_ becomes excessively large, the off-diagonal entries *K*_*ij*_ can approach zero, causing the kernel matrix to resemble the identity. Because *∂K*_*ij*_*/∂r*_*m*_ = *−*(*z*_*im*_*−z*_*jm*_)^2^*K*_*ij*_, the corresponding derivatives also vanish, making the likelihood increasingly insensitive to further changes in *r*_*m*_. Consequently, distinguishing feature relevance becomes more difficult, particularly for modest sample sizes *N* . This pattern is evident in Figure 2, where most methods attain their highest AUPRC at intermediate values of *r*_*m*_. RKMR outperformed all competing methods across nearly all settings; at the largest value of *r*_*m*_, RF marginally outperformed RKMR. RF was otherwise the closest competitor, although its average AUPRC remained at least 30% lower. Enet also performed reasonably well at smaller values of *r*_*m*_, whereas the specialized marker-selection methods were consistently outperformed by both RF and Enet across all parameter settings.

**Figure 2:**
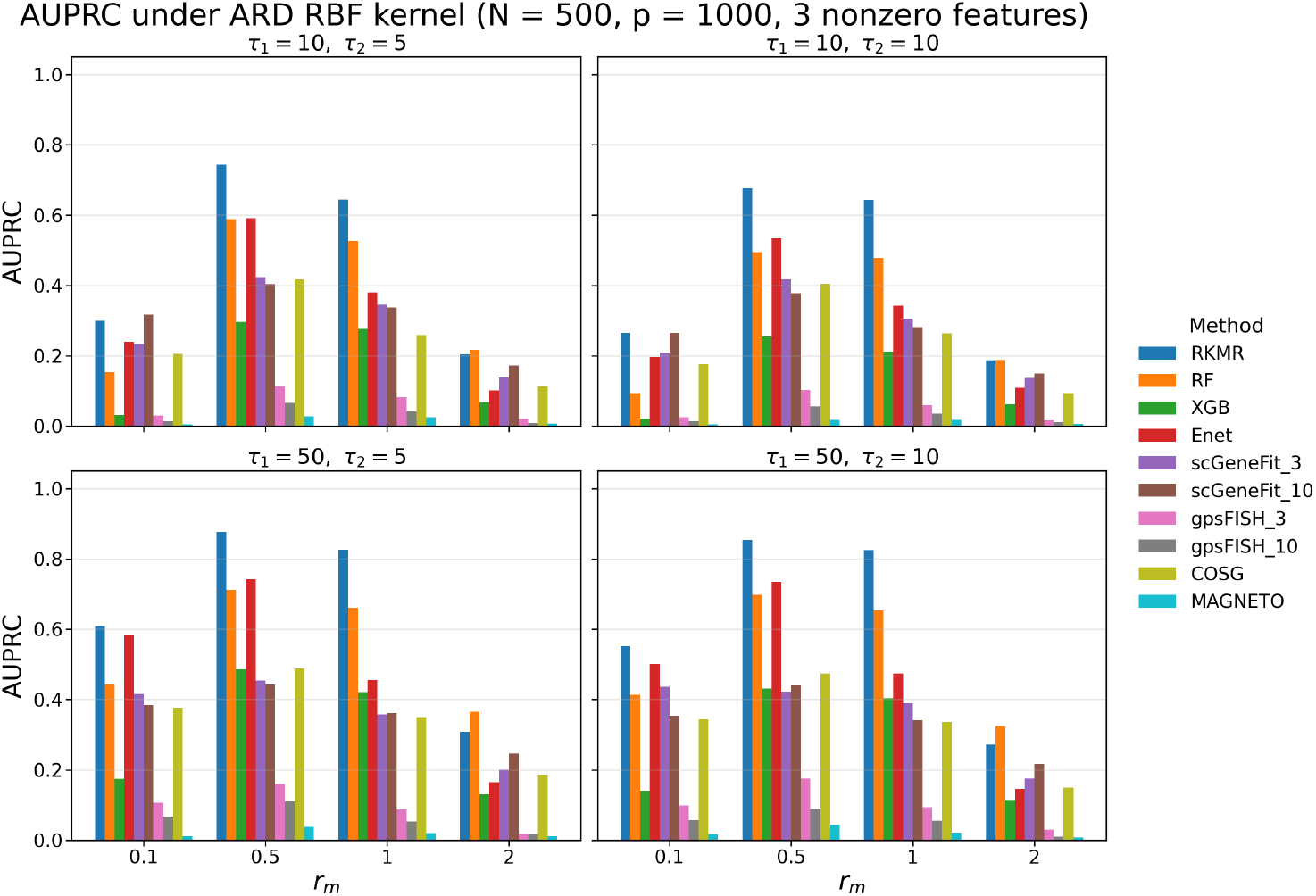
AUPRC comparisons for Simulation Scenario 2 across parameter settings. Increasing *r*_*m*_ strengthens feature relevance, but excessively large values can make the ARD RBF kernel matrix approach the identity matrix and hence, hinder feature recovery.

Overall, RKMR showed the strongest and most consistent feature-recovery performance across both scenarios. Enet remained competitive under linear effects, whereas RF was the strongest alternative under the nonlinear ARD RBF kernel. Performance generally declined as the spatial effect increased and, under the RBF kernel, often peaked at intermediate values of *r*_*m*_. Although AUPRC may appear favorable for some methods, RKMR additionally provides probabilistic selection through PIPs, offering a more informative measure of biological relevance.

## 4 Real data analysis

A key advantage of RKMR is its applicability to both spatial and nonspatial single-cell data within a unified framework: for scRNA-seq datasets, the spatial kernel can simply be omitted without altering the feature-selection machinery. To demonstrate this flexibility, we evaluated RKMR on two scRNA-seq datasets and two ST datasets.

### 4.1 Pancreatic islet cells data

Lawlor et al. (2017) ^91^ generated single-cell transcriptomes measuring 26,616 genes across 638 human pancreatic islet cells from non-diabetic and type 2 diabetic (T2D) donors. Their analysis characterized transcriptional signatures of the major endocrine islet cell types, including *alpha, beta, delta*, and *Gamma/PP* cells, and identified cell-type-specific expression changes in T2D that were not detectable through bulk islet profiling. The reported canonical markers included INS (*beta*), GCG (*alpha*), SST (*delta*), PPY (*gamma/PP*), PRSS1 (*acinar*), COL1A1 (*stellate*), and KRT19 (*ductal*). After standard quality-control and filtering procedures, we retained 604 cells and the top 3,000 highly variable genes (HVGs). To reduce multicollinearity, we applied correlation-based filtering using the `caret` package in `R`, removing genes until all pairwise correlations were less than or equal to 0.5. Because this step excluded some reported markers that were highly correlated with other genes, we manually restored the top reported marker for each cell type, yielding 2,922 genes for analysis. As shown in Table 2, RKMR selected exactly one gene with PIP *>* 0.5 for each cell type, and in every case it was the reported canonical marker, demonstrating highly specific marker recovery.

**Table 2:**
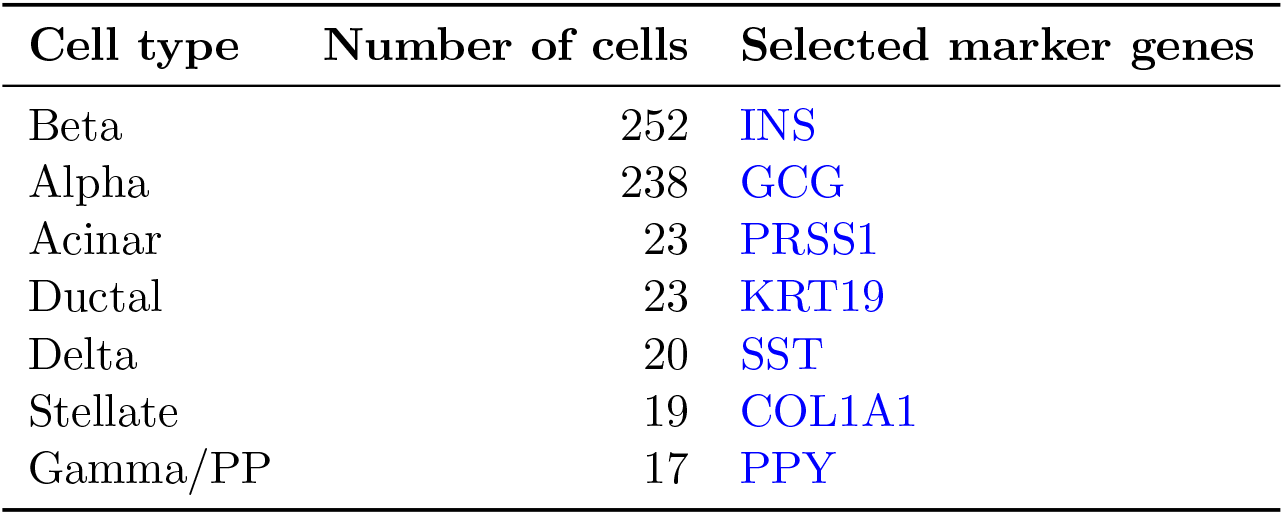
Cell-type frequencies and RKMR-selected markers with PIP *>* 0.5. Blue denotes reported markers from Lawlor et al. (2017), all uniquely recovered by RKMR.

| Cell type | Number of cells | Selected marker genes |
| --- | --- | --- |
| Beta | 252 | INS |
| Alpha | 238 | GCG |
| Acinar | 23 | PRSS1 |
| Ductal | 23 | KRT19 |
| Delta | 20 | SST |
| Stellate | 19 | COL1A1 |
| Gamma/PP | 17 | PPY |

### 4.2 Mouse cortex scRNA-seq data

We next analyzed a mouse cortex and hippocampus scRNA-seq dataset comprising 20,006 genes measured across 3,005 cells from nine cell types ^92^. Zeisel et al. (2015) used large-scale scRNA-seq to establish a transcriptomic taxonomy of cortical and hippocampal cell types and identify marker genes linking these cell types to known anatomical and cellular classes. Following the preprocessing and correlation-filtering protocol described above, we reduced the 3,000 most highly variable genes to 1,256 genes, while manually restoring any reported markers removed because of correlations exceeding 0.5. As shown in Table 3, RKMR recovered the reported marker for every cell type except *Microglia*, for which the reported marker was Aif1. Instead, RKMR selected the correlated gene Lyz2 (*r* = 0.63), which is biologically plausible because Lyz2 and Aif1 are associated with related microglial and myeloid expression programs ^93^. For several cell types, including *Endothelial* and *Mural*, RKMR selected no more than two genes while retaining the reported marker, demonstrating its ability to identify highly specific marker panels. RKMR also recovered the reported marker for the rare *Ependymal* population, which comprised only 30 cells (1% of the dataset).

**Table 3:**
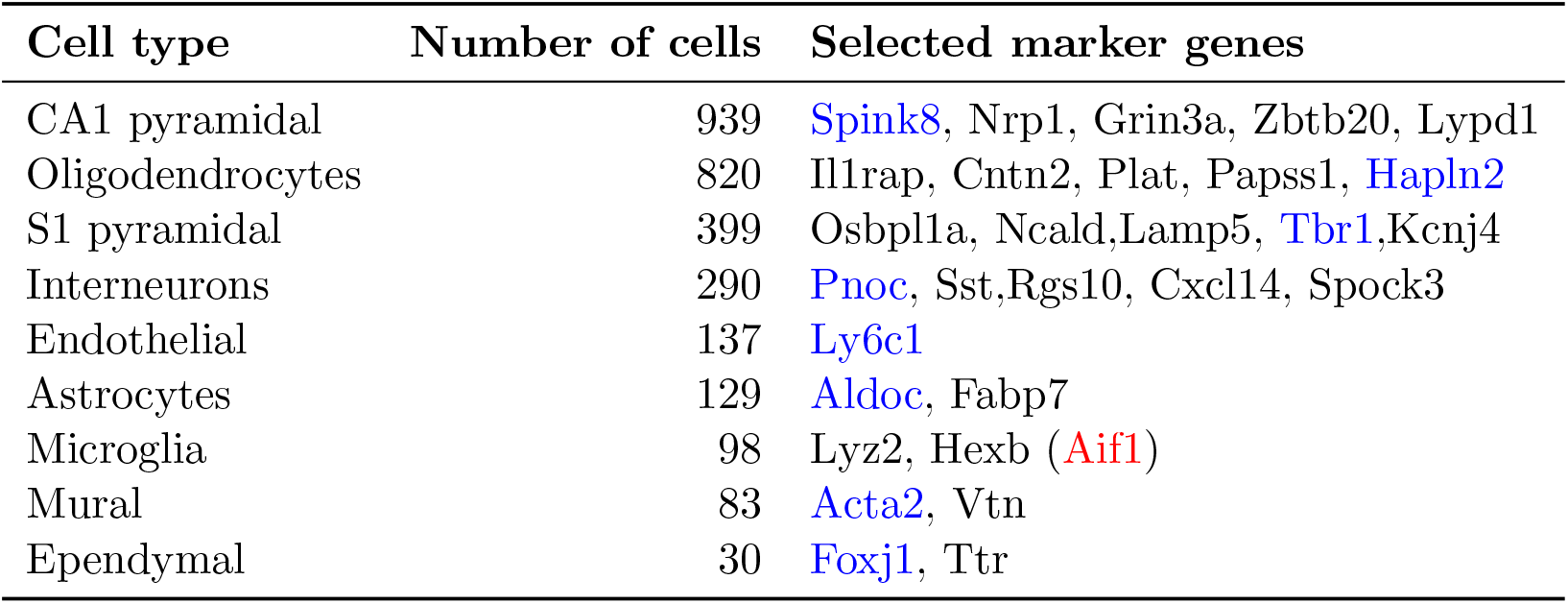
Cell-type frequencies and the top five RKMR-selected marker genes with PIP *>* 0.5. Blue denotes top markers reported by Zeisel et al. (2015) that were recovered by RKMR, whereas red denotes a reported marker that was not selected.

### 4.3 Axolotl brain ST data

We next analyzed a single-cell-resolution Stereo-seq dataset of axolotl brain development and regeneration ^94^. Wei et al. (2022) generated spatial transcriptomic maps across developmental and regenerative stages and identified an injury-induced ependymoglial progenitor population potentially involved in replenishing lost neurons. We focused on the Stage 57 section, comprising 22,893 genes measured across 4,410 cells from 17 spatially organized cell types (Fig. 3A), each with a reported top marker gene. Following the preprocessing and correlation-filtering protocol described above, we reduced the 3,000 most highly variable genes to 2,221 genes, while manually restoring reported markers.

**Figure 3:**
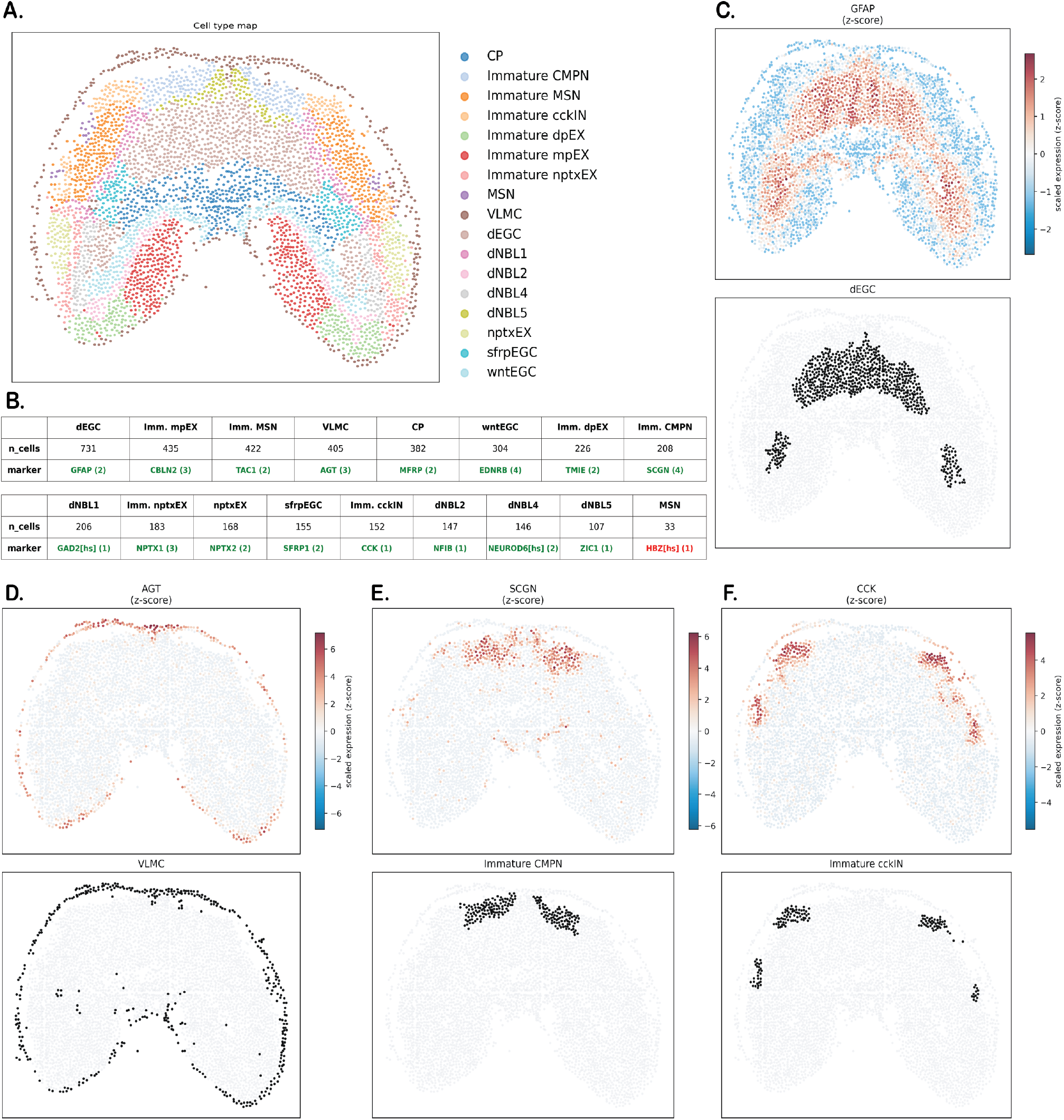
**A**. Spatial organization of the 17 cell types. **B**. Number of cells per cell type and the top marker gene reported by Wei et al. (2022). Green indicates that RKMR identified the reported marker, whereas red indicates that it was not selected. The number beside each marker denotes the size of the marker panel selected by RKMR (PIP *>* 0.5). **C–F**. Spatial expression patterns of GFAP in *dEGC* cells, AGT in *VLMC* cells, SCGN in *Immature CMPN* cells, and CCK in *Immature cckIN* cells. The corresponding cell-type organization is shown individually in the bottom panels.

Figure 3B shows the number of cells (*n* cells) and the top reported marker for each cell type. Using a PIP threshold of 0.5, RKMR identified all reported markers except HBZ[hs] for MSN, for which it selected the correlated alternative HBG2[hs] (*r* = 0.67). Because both genes are involved in related hemoglobin-associated erythroid functions, the selection of HBG2[hs] is biologically supported ^95^. For every other cell type, the reported marker was the highest-ranked selected gene, with a PIP of 1. RKMR selected only one to four genes per cell type, demonstrating its ability to recover highly sparse marker sets. Figures 3C–F display the expression patterns of the reported and RKMR-selected markers for *dEGC* (731 cells), *VLMC* (405 cells), *Immature CMPN* (208 cells), and *Immature cckIN* (152 cells), illustrating effective marker recovery across both abundant and relatively rare cell populations. The spatial organization of these cell types also reveals substantial spatial complexity in the outcome patterns, underscoring the importance of explicitly accounting for spatial dependence, as incorporated in RKMR.

### 4.4 DLPFC data

Finally, we analyzed the human dorsolateral prefrontal cortex (DLPFC) Visium dataset, comprising 12 tissue sections from three donors^96^. Maynard et al. (2021) generated transcriptome-wide spatial maps of the six cortical layers and white matter (WM), identifying extensive layer-enriched expression patterns in the adult human DLPFC. Each section contains approximately 4,000 spots profiled for 33,538 genes, with spot-level layer annotations (Fig. 4). Following a preprocessing and filtering protocol similar to that described above, we retained a set of 3,000 HVGs across sections. Although Maynard et al. (2021) evaluated previously reported markers and identified several novel layer-enriched genes, they did not define a single curated marker panel for each layer. We therefore applied RKMR separately to each section to identify layer-specific markers and assess the reproducibility of these discoveries across sections.

**Figure 4:**
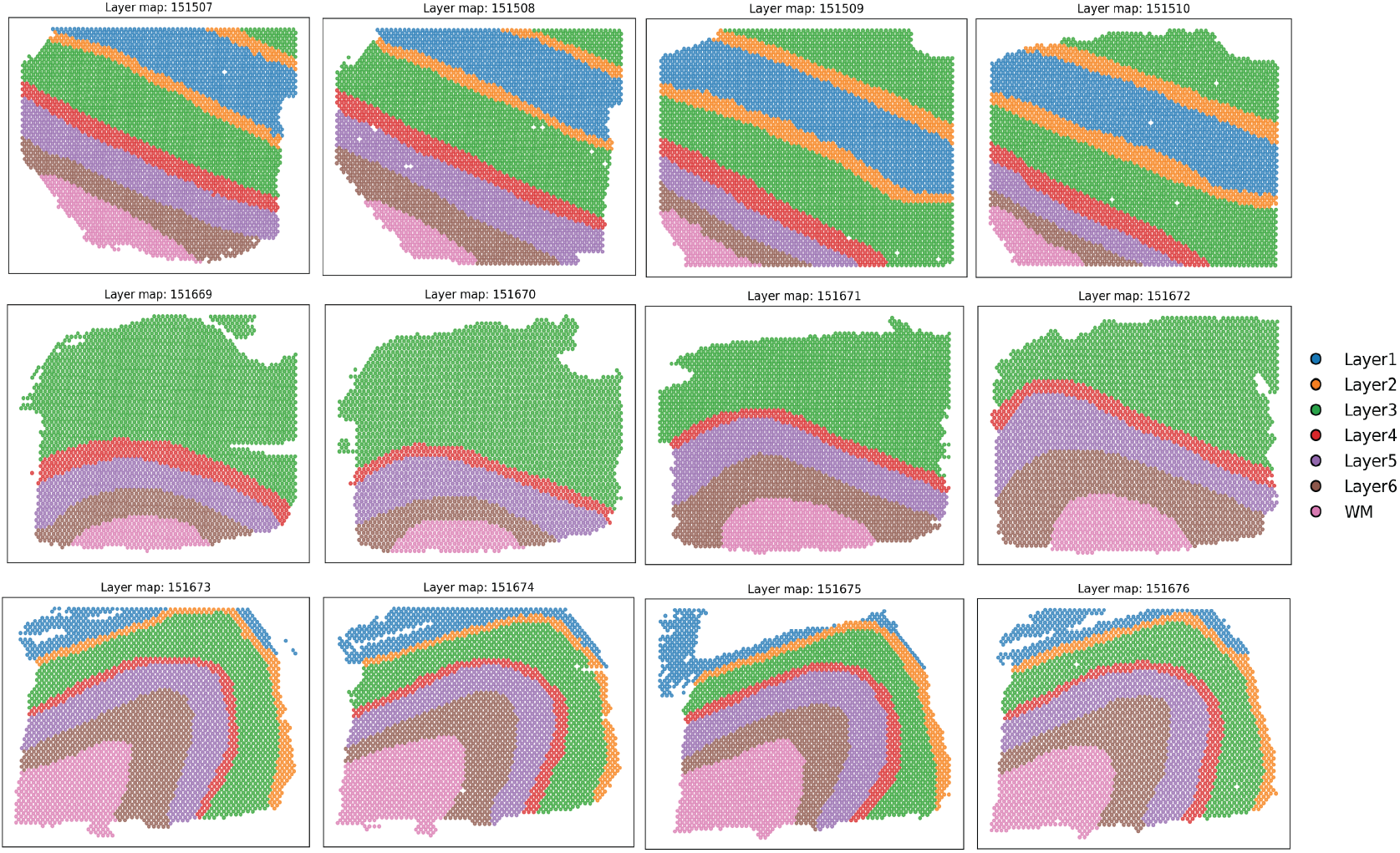
Manual spot-level annotations of cortical layers and white matter across the 12 DLPFC sections from Maynard et al. (2021) ^96^. Sections 151669–151672 do not contain Layers 1 and 2.

Table 4 shows that several genes were identified for the same layer in more than four sections, as highlighted in blue. Most were either reported by Maynard et al. (2021) or have prior support as layer-associated canonical markers. Representative examples include GFAP for *Layer 1* ^97^, CAMK2N1 for *Layer 2* ^98^, PVALB for *Layer 4* ^99^, PCP4 for *Layer 5* ^100^, CCK for *Layer 6* ^101^, and MBP for *WM* ^102^. Some markers, particularly those identified for *Layers 2 and 3*, may not be uniquely restricted to a single layer. Because RKMR performs *one-versus-rest* marker selection separately for each layer, genes enriched in adjacent or transcriptionally similar layers may be selected for both. Figure 5 further illustrates the difficulty of layer-specific marker recovery, as the expression of the representative markers PCP4 and CCK is not always sharply confined to the corresponding layer boundaries and includes substantial background expression throughout the tissue sections. Nevertheless, RKMR consistently recovered these markers, demonstrating its robustness to the noisy expression patterns commonly encountered in real ST data.

**Table 4:**
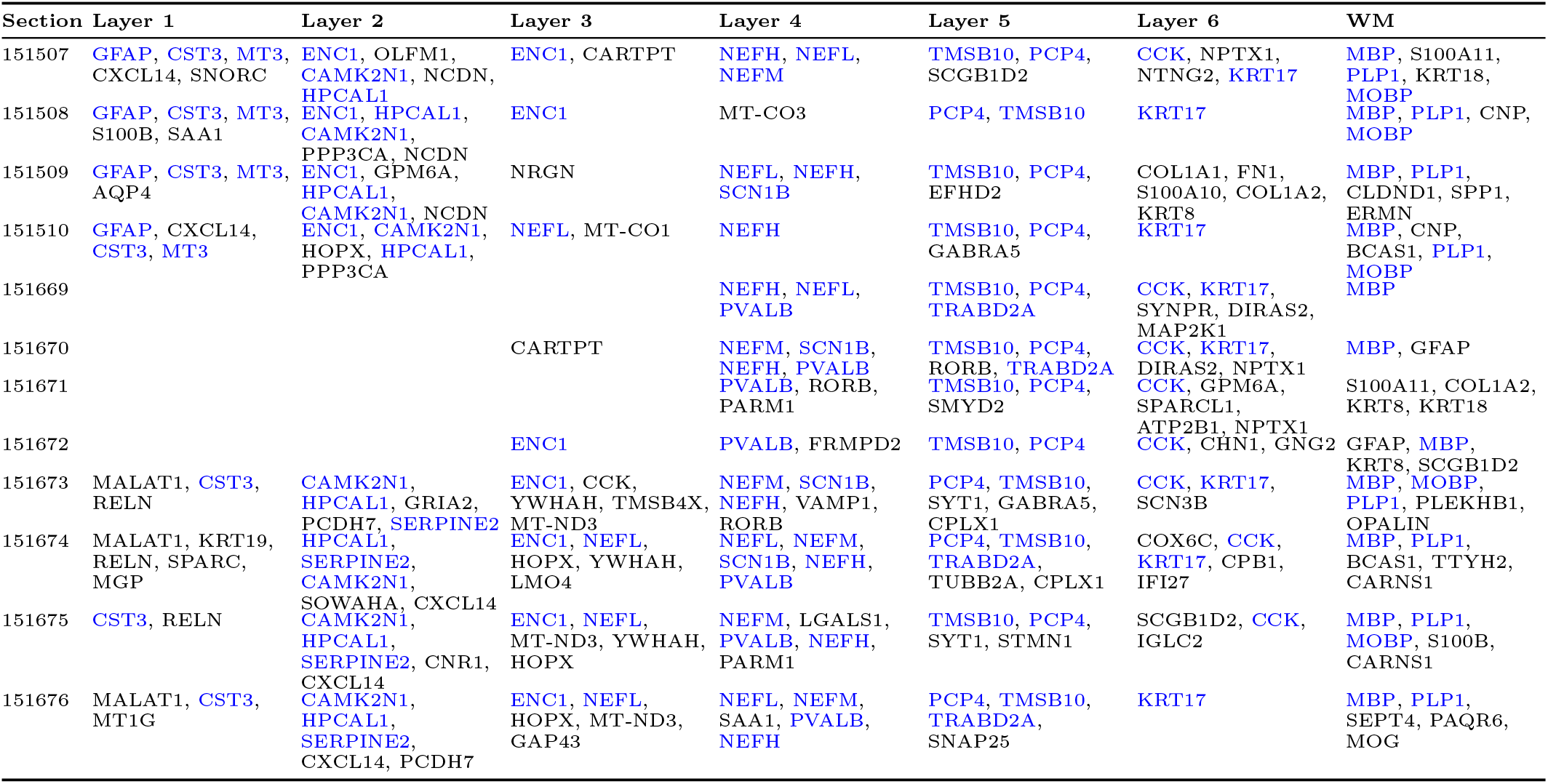
Selected top five marker genes with PIP *>* 0.5 by DLPFC section and layer. Genes appearing in at least four sections within the same layer are shown in blue.

**Figure 5:**
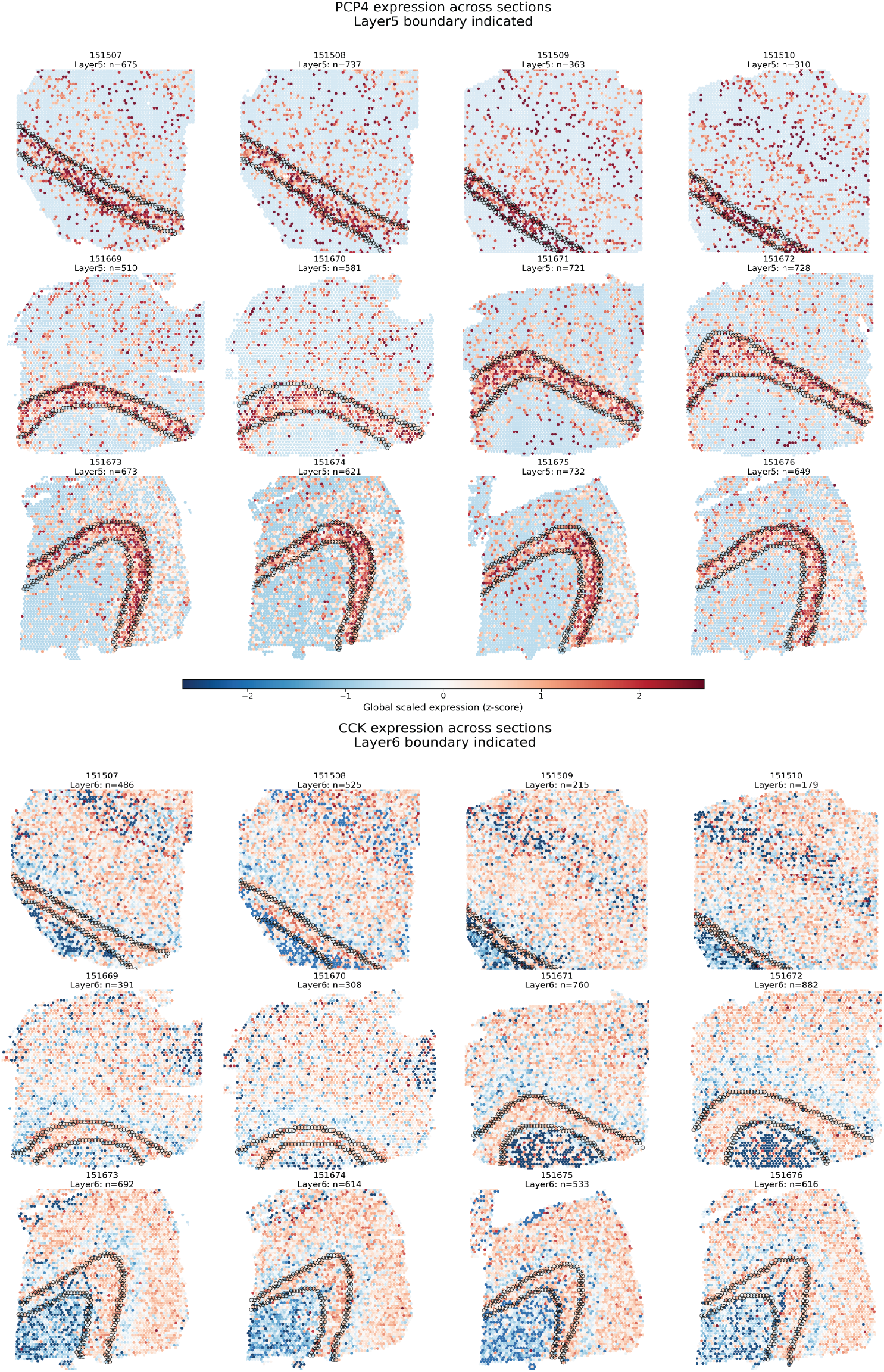
Expression of the detected markers PCP4 and CCK for *Layers 5 and 6*, respectively, across tissue sections. Layer boundaries are outlined in black.

## 5 Conclusion

Marker-panel discovery—the identification of a small set of discriminative and potentially actionable features for distinguishing known cell types, tissue layers, or disease subclasses—remains understudied in spatial transcriptomics and spatial proteomics. Existing approaches often rely on methods originally developed for scRNA-seq data, many of which neither explicitly account for tissue spatial structure nor provide probabilistic selection criteria. To address this gap, we introduce rapid kernel machine regression (RKMR), a flexible framework that jointly models nonlinear feature–outcome relationships and spatial dependence through kernel-based components. By combining an automatic relevance determination (ARD) kernel with a semi-Bayesian spike-and-slab formulation, RKMR provides interpretable feature selection through posterior inclusion probabilities (PIPs). RKMR further integrates variational inference with low-rank kernel approximations to improve computational scalability, facilitating its application to increasingly large spatial omics datasets.

Across extensive simulation studies spanning different levels of feature effects and spatial autocorrelation in the outcome, RKMR outperformed not only general-purpose algorithms such as random forest, XG-Boost, and elastic net, but also several specialized marker-discovery methods developed for single-cell data. For most competing methods, the final selection rule is either user-specified or based on data-dependent importance scores. In contrast, RKMR provides a probabilistic and readily interpretable criterion based on PIPs, such as PIP *>* 0.5. RKMR can also be applied to scRNA-seq data by omitting the spatial kernel. In analyses of pancreatic islet cells and mouse cortex scRNA-seq data ^91,92^, RKMR recovered the leading marker genes reported in the original studies while automatically producing sparse panels that typically contained the top marker and, in some cases, a small number of correlated genes. Similarly, in a Stereo-seq dataset of axolotl brain regeneration ^94^, RKMR recovered the top reported marker for 16 of 17 cell types. In the Visium DLPFC dataset comprising 12 tissue sections ^96^, RKMR identified largely consistent marker sets across sections for each of the six cortical layers and white matter, demonstrating its robustness and reproducibility in noisy real-world datasets.

A persistent challenge in marker discovery is multicollinearity arising from gene co-expression ^103^, which can lead to the selection of a correlated feature as a proxy for a known marker, as observed in several of our real-data analyses. We briefly explored correlation-aware priors for the spike-and-slab inclusion indicators^104,105^, but they did not consistently improve performance in the present setting. Developing structured priors or joint panel-selection strategies that more effectively account for correlated and redundant features therefore represents an important direction for future work. RKMR currently uses a *one-versus-rest* formulation, analyzing each target cell type or tissue class against all remaining observations. A natural extension would replace the binary logistic model with a multinomial logistic formulation, enabling joint marker selection across multiple classes. The increasing availability of spatial multi-omics data also motivates extensions to multimodal marker discovery. Such models must carefully accommodate differences in measurement scale, modality-specific noise, and cross-modal dependence ^106^. To facilitate broader use, we have developed a computationally efficient `Python` implementation with user-oriented documentation and tutorials. The software will be made publicly available at https://github.com/sealx017/RKMR.

