## Supplementary Material for "RKMR: A Rapid Kernel Machine Regression Framework for Optimal Marker Detection in Spatial Omics Data"

August 3, 2026

### 1 Full algorithmic details

Notations from the main paper remain unchanged. Let $\mathbf{H} = [\mathbf{1}_N\ \mathbf{X}]$ denote the fixed-effect design matrix and $\boldsymbol{\beta} = (\beta_0, \boldsymbol{\beta}_1^\top)^\top$ the corresponding coefficient vector. For an intercept-only model, set $\mathbf{H} = \mathbf{1}_N$. Let

$\mathcal{L} \subset \{1, \ldots, N\}$ denote the set of Nyström landmarks, with $|\mathcal{L}| = L$, kept at 500. For the full-data PQL/IRLS update, define the combined cross-kernel matrix, landmark kernel matrix, and Nyström approximation by

$$\begin{aligned} \mathbf{C} &= \tau_1 K_{\mathbf{r}}(\mathbf{Z}, \mathbf{Z}_{\mathcal{L}}) + \tau_2 C_\rho(\mathbf{S}, \mathbf{S}_{\mathcal{L}}), \\ \mathbf{K}_{\mathcal{L}\mathcal{L}} &= \tau_1 K_{\mathbf{r}}(\mathbf{Z}_{\mathcal{L}}, \mathbf{Z}_{\mathcal{L}}) + \tau_2 C_\rho(\mathbf{S}_{\mathcal{L}}, \mathbf{S}_{\mathcal{L}}) + \epsilon_{\text{nys}} \mathbf{I}_L, \\ \mathbf{\Phi} &= \mathbf{C}\mathbf{K}_{\mathcal{L}\mathcal{L}}^{-1/2}, \qquad \mathbf{Q} = \mathbf{\Phi}\mathbf{\Phi}^\top = \mathbf{C}\mathbf{K}_{\mathcal{L}\mathcal{L}}^{-1}\mathbf{C}^\top. \end{aligned} \tag{1}$$

Thus, the Nyström approximation is applied only to the full combined molecular–spatial kernel matrix used in the PQL/IRLS step. Minibatching is a separate approximation used only to form stochastic gradients of the Gaussian-surrogate objective. In the algorithms, $r_m$ denotes the zero-temperature point-mass location (the $\mu_m$ of the main text), so $\mathbb{E}_{q(\mathbf{r})}(r_m^2) = r_m^2$.

For a sampled minibatch $\mathcal{B} \subset \{1, \ldots, N\}$, define the exact combined minibatch kernel and working covariance as

$$\begin{aligned} \mathbf{K}_{\mathcal{B}} &= \tau_1 K_{\mathbf{r}}(\mathbf{Z}_{\mathcal{B}}, \mathbf{Z}_{\mathcal{B}}) + \tau_2 C_\rho(\mathbf{S}_{\mathcal{B}}, \mathbf{S}_{\mathcal{B}}), \\ \tilde{\mathbf{K}}_{\mathcal{B}} &= \mathbf{K}_{\mathcal{B}} + \operatorname{diag}(1/\mathbf{w}_{\mathcal{B}}). \end{aligned} \tag{2}$$

No Nyström approximation is applied to $\mathbf{K}_{\mathcal{B}}$. The required minibatch kernel derivatives are

$$\begin{aligned} \mathbf{G}_{\mathcal{B}}^{(r_m)} &= \tau_1 \frac{\partial K_{\mathbf{r}}(\mathbf{Z}_{\mathcal{B}}, \mathbf{Z}_{\mathcal{B}})}{\partial r_m}, \quad & \mathbf{G}_{\mathcal{B}}^{(\rho)} &= \tau_2 \frac{\partial C_\rho(\mathbf{S}_{\mathcal{B}}, \mathbf{S}_{\mathcal{B}})}{\partial \rho}, \\ \mathbf{G}_{\mathcal{B}}^{(\tau_1)} &= K_{\mathbf{r}}(\mathbf{Z}_{\mathcal{B}}, \mathbf{Z}_{\mathcal{B}}), & \mathbf{G}_{\mathcal{B}}^{(\tau_2)} &= C_\rho(\mathbf{S}_{\mathcal{B}}, \mathbf{S}_{\mathcal{B}}). \end{aligned} \tag{3}$$

**Algorithm 1** Nyström-accelerated stochastic a-CAVI for logistic RKMR

**Require:** $\mathbf{y} \in \{0,1\}^N$, $\mathbf{Z} \in \mathbb{R}^{N \times p}$, $\mathbf{S}$, $\mathbf{X}$; kernels $K_{\mathbf{r}}$ and $C_\rho$
**Require:** Hyperparameters $(v, c, a, b)$; minibatch size $B$; landmarks $L$; outer iterations $T$; inner steps $\{G_t\}_{t=1}^T$; ridge $\epsilon_{\text{nys}} > 0$
Set $\mathbf{H} \leftarrow [\mathbf{1}_N \ \mathbf{X}]$; select $\mathcal{L} \subset \{1, \ldots, N\}$, $|\mathcal{L}| = L$; form $\mathbf{Z}_{\mathcal{L}}$ and $\mathbf{S}_{\mathcal{L}}$
Initialize $\mathbf{r}, \rho, \tau_1, \tau_2, \boldsymbol{\lambda}, \boldsymbol{\beta}, \boldsymbol{\eta}, \xi_a, \xi_b$
Compute $\mathbb{E}_{q(\pi)}[\log \pi]$ and $\mathbb{E}_{q(\pi)}[\log(1-\pi)]$
**for** $t = 1, \ldots, T$ **do**
  Update $(\boldsymbol{\beta}, \boldsymbol{\eta}, \boldsymbol{\mu}, \mathbf{w}, \tilde{\mathbf{y}})$ using Algorithm 2
  Update $(\mathbf{r}, \rho, \tau_1, \tau_2, \boldsymbol{\lambda}, \xi_a, \xi_b, \mathbb{E}_{q(\pi)}[\log \pi], \mathbb{E}_{q(\pi)}[\log(1-\pi)])$ using Algorithm 3
  **if** the model and variational parameters have converged **then**
    **break**
  **end if**
**end for**
Run Algorithm 2 once more at the final $(\mathbf{r}, \rho, \tau_1, \tau_2)$
**return** $(\mathbf{r}, \rho, \tau_1, \tau_2, \boldsymbol{\lambda}, \beta_0, \boldsymbol{\beta}_1, \boldsymbol{\eta})$

**Algorithm 2** Nyström-accelerated IRLS update for logistic RKMR

**Require:** $\mathbf{y}, \mathbf{Z}, \mathbf{S}, \mathbf{H}, \mathbf{Z}_{\mathcal{L}}, \mathbf{S}_{\mathcal{L}}; \mathbf{r}, \rho, \tau_1, \tau_2, \boldsymbol{\eta}, \epsilon_{\text{nys}}$

$\mathbf{C} \leftarrow \tau_1 K_{\mathbf{r}}(\mathbf{Z}, \mathbf{Z}_{\mathcal{L}}) + \tau_2 C_\rho(\mathbf{S}, \mathbf{S}_{\mathcal{L}})$

$\mathbf{K}_{\mathcal{L}\mathcal{L}} \leftarrow \tau_1 K_{\mathbf{r}}(\mathbf{Z}_{\mathcal{L}}, \mathbf{Z}_{\mathcal{L}}) + \tau_2 C_\rho(\mathbf{S}_{\mathcal{L}}, \mathbf{S}_{\mathcal{L}}) + \epsilon_{\text{nys}} \mathbf{I}_L$

$\boldsymbol{\Phi} \leftarrow \mathbf{C}\mathbf{K}_{\mathcal{L}\mathcal{L}}^{-1/2}$; $\mathbf{Q} \leftarrow \boldsymbol{\Phi}\boldsymbol{\Phi}^\top \approx \tau_1 \mathbf{K}_{\mathbf{r}} + \tau_2 \mathbf{C}_\rho$

$\boldsymbol{\mu} \leftarrow \sigma(\boldsymbol{\eta})$; $w_i \leftarrow \mu_i(1-\mu_i)$; $\tilde{y}_i \leftarrow \eta_i + (y_i - \mu_i)/w_i$, $i = 1, \ldots, N$

$\mathbf{D} \leftarrow \text{diag}(1/\mathbf{w}) = \mathbf{W}^{-1}$

Define $\mathcal{S}(\mathbf{U}) = (\mathbf{Q} + \mathbf{D})^{-1}\mathbf{U}$ using Woodbury:

$\quad \mathcal{S}(\mathbf{U}) = \mathbf{D}^{-1}\mathbf{U} - \mathbf{D}^{-1}\boldsymbol{\Phi}(\mathbf{I}_L + \boldsymbol{\Phi}^\top \mathbf{D}^{-1}\boldsymbol{\Phi})^{-1}\boldsymbol{\Phi}^\top \mathbf{D}^{-1}\mathbf{U}.$

$\mathbf{z}_y \leftarrow \mathcal{S}(\tilde{\mathbf{y}})$; $\mathbf{Z}_H \leftarrow \mathcal{S}(\mathbf{H})$

$\boldsymbol{\beta} \leftarrow (\mathbf{H}^\top \mathbf{Z}_H)^{-1}\mathbf{H}^\top \mathbf{z}_y$, where $\boldsymbol{\beta} = (\beta_0, \boldsymbol{\beta}_1^\top)^\top$

$\boldsymbol{\alpha} \leftarrow \mathcal{S}(\tilde{\mathbf{y}} - \mathbf{H}\boldsymbol{\beta})$

$\boldsymbol{\eta} \leftarrow \mathbf{H}\boldsymbol{\beta} + \boldsymbol{\Phi}(\boldsymbol{\Phi}^\top \boldsymbol{\alpha})$

Recompute $\boldsymbol{\mu}, \mathbf{w}, \tilde{\mathbf{y}}$ using the updated $\boldsymbol{\eta}$

**return** $(\boldsymbol{\beta}, \boldsymbol{\eta}, \boldsymbol{\mu}, \mathbf{w}, \tilde{\mathbf{y}})$

**Algorithm 3** Stochastic ARD, kernel-parameter, and inclusion-probability updates

**Require:** $\tilde{\mathbf{y}}, \mathbf{w}, \mathbf{Z}, \mathbf{S}, \mathbf{H}; \mathbf{r}, \rho, \tau_1, \tau_2, \boldsymbol{\lambda}$

**Require:** $\mathbb{E}_{q(\pi)}[\log \pi], \mathbb{E}_{q(\pi)}[\log(1-\pi)]$; $(v, c, a, b)$; minibatch size $B$; inner steps $G_t$

**for** $g = 1, \ldots, G_t$ **do**

  Sample $\mathcal{B} \subset \{1, \ldots, N\}$ with $|\mathcal{B}| = B$

  Compute the exact combined minibatch kernel $\mathbf{K}_{\mathcal{B}}$ using Eq. (2)

  $\tilde{\mathbf{K}}_{\mathcal{B}} \leftarrow \mathbf{K}_{\mathcal{B}} + \text{diag}(1/\mathbf{w}_{\mathcal{B}})$; $\mathbf{V}_{\mathcal{B}} \leftarrow \tilde{\mathbf{K}}_{\mathcal{B}}^{-1}$

  $\hat{\boldsymbol{\beta}}_{\mathcal{B}} \leftarrow (\mathbf{H}_{\mathcal{B}}^\top \mathbf{V}_{\mathcal{B}} \mathbf{H}_{\mathcal{B}})^{-1}\mathbf{H}_{\mathcal{B}}^\top \mathbf{V}_{\mathcal{B}} \tilde{\mathbf{y}}_{\mathcal{B}}$

  $\mathbf{e}_{\mathcal{B}} \leftarrow \tilde{\mathbf{y}}_{\mathcal{B}} - \mathbf{H}_{\mathcal{B}}\hat{\boldsymbol{\beta}}_{\mathcal{B}}$

  $\mathbf{A}_{\mathcal{B}} \leftarrow \mathbf{V}_{\mathcal{B}}\mathbf{e}_{\mathcal{B}}\mathbf{e}_{\mathcal{B}}^\top \mathbf{V}_{\mathcal{B}} + \mathbf{V}_{\mathcal{B}}\mathbf{H}_{\mathcal{B}}(\mathbf{H}_{\mathcal{B}}^\top \mathbf{V}_{\mathcal{B}}\mathbf{H}_{\mathcal{B}})^{-1}\mathbf{H}_{\mathcal{B}}^\top \mathbf{V}_{\mathcal{B}} - \mathbf{V}_{\mathcal{B}}.$

  **for** $m = 1, \ldots, p$ **do**

    Compute $\mathbf{G}_{\mathcal{B}}^{(r_m)}$ using Eq. (3)

    $\hat{\nabla}_{r_m} \leftarrow \dfrac{N}{2B}\langle \mathbf{A}_{\mathcal{B}}, \mathbf{G}_{\mathcal{B}}^{(r_m)} \rangle_F - v\{(1-\lambda_m) + c\lambda_m\} r_m$

  **end for**

  **for** $\theta \in \{\rho, \tau_1, \tau_2\}$ **do**

    Compute $\mathbf{G}_{\mathcal{B}}^{(\theta)}$ using Eq. (3)

    $\hat{\nabla}_\theta \leftarrow \dfrac{N}{2B}\langle \mathbf{A}_{\mathcal{B}}, \mathbf{G}_{\mathcal{B}}^{(\theta)} \rangle_F$

  **end for**

  Update $(\mathbf{r}, \rho, \tau_1, \tau_2)$ by Adam or stochastic gradient ascent, enforcing $\rho, \tau_1, \tau_2 > 0$

**end for**

**for** $m = 1, \ldots, p$ **do**

  Set $\mathbb{E}_{q(\mathbf{r})}(r_m^2) \leftarrow r_m^2$ under the zero-temperature approximation

  $\lambda_m \leftarrow \left[1 + c^{-1/2}\exp\left\{-\dfrac{1}{2}v(1-c)r_m^2 + \mathbb{E}_{q(\pi)}[\log(1-\pi)] - \mathbb{E}_{q(\pi)}[\log \pi]\right\}\right]^{-1}$

**end for**

$\xi_a \leftarrow a + \sum_{m=1}^p \lambda_m$; $\xi_b \leftarrow b + p - \sum_{m=1}^p \lambda_m$

$\mathbb{E}_{q(\pi)}[\log \pi] \leftarrow \psi(\xi_a) - \psi(\xi_a + \xi_b)$; $\mathbb{E}_{q(\pi)}[\log(1-\pi)] \leftarrow \psi(\xi_b) - \psi(\xi_a + \xi_b)$

**return** $(\mathbf{r}, \rho, \tau_1, \tau_2, \boldsymbol{\lambda}, \xi_a, \xi_b, \mathbb{E}_{q(\pi)}[\log \pi], \mathbb{E}_{q(\pi)}[\log(1-\pi)])$
